# Identification of key immune regulatory genes in HIV–1 Progression

**DOI:** 10.1101/2020.10.09.333716

**Authors:** Sk Md Mosaddek Hossain, Lutfunnesa Khatun, Sumanta Ray, Anirban Mukhopadhyay

## Abstract

In the last few decades, application of DNA microarray technology has sprung up as a powerful technique for discovering stage specific changes in expression pattern of a disease progression. Human Immunodeficiency Virus (HIV) infection causes Acquired Immunodeficiency Syndrome (AIDS) which is one of the most devastating diseases affecting humankind. Here, we have proposed a framework to examine the difference among microarray gene expression data of uninfected and three different HIV–1 infection stages using module preservation statistics. Initially, we detected differentially expressed genes among all the stages and identified coexpression modules by using topological overlap as a dissimilarity measure. To examine relationship among co-expression modules, we have compiled a module eigenegene network for each sample category which models similarity among all coexpression modules. To further examine the network, we have found clusters in it which are termed as ‘meta-modules’. Different module preservation statistics with two composite statistics: “*Z*_*summary*_” and “*MedianRank*” are utilized to examine changes in structure of coexpression modules. We have applied our proposed methodology to discover modular changes between uninfected and acute samples, acute and chronic samples, chronic and AIDS samples. We have found several interesting results on preservation characteristics of gene modules across different stages. Some genes are identified to be preserved in a pair of stages while alter their characteristics across other stages. We further validated the obtained results using permutation test and classification techniques. Biological significance of the obtained modules have been examined using gene ontology and pathway based analysis. Additionally, we have detected key immune regulatory hub genes in the associated protein-protein interaction networks (PPINs) of the differentially expressed genes (DEGs) using twelve topological and centrality analysis methods. Moreover, we have analyzed the key immune regulatory genes which interacts with HIV-1 proteins inside the preserved and perturbed meta-modules across different HIV-1 stages and thus likely to act as potential biomarkers in HIV–1 progression.

## 1 Introduction

Human Immunodeficiency Virus (HIV) is a type of retrovirus that attacks the immune system of the body and in almost all instances finally leads to acquired immunodeficiency syndrome (AIDS). AIDS is a collection of several diseases and infections. HIV infection occurs as a consequence of contamination with the HIV virus via body fluids from an already infected person. HIV-1 is the most common, predominant type among the two main HIV types: HIV-1, HIV-2. The health of the HIV affected people depends on the condition of their immune system, specially on the count of CD4+ T cells and number of viral particles present in the blood (i.e. viral load). HIV-infection cannot incisively demarked into distinct phases, although typically there are 3 different stages in which HIV prolongs: acute, chronic and AIDS.

The acute phase begins immediately after someone is infected with HIV and remains till the seroconversion takes place. Acute phase typically lasts roughly about 2–4 weeks, in this period HIV RNA levels and the plasma p24 antigen are discernible [1]. At this stage, HIV virus replicates itself rapidly in the body and destroys the infection fighting CD4+ T cells making HIV infected persons more vulnerable to opportunistic infections [2], [3]. Most of the people at this stage perceives flu-like symptoms including fever, headache, sore throat, rash, muscle and joint pains which exemplify the body’s natural reaction to an infection as it attempts to eradicate the virus. Some people may not experience any symptoms at all in this stage [4].

After the acute stage, HIV-1 infected persons proceed into a long term phase known as chronic or clinical latency stage. At this stage HIV virus goes on replicating itself but at a very low rate and continues to destroy victims immune systems as CD4+ T cell counts gradually decreases. People generally experience no symptom (asymptotic) or only mild ones (symptomatic) and thus this stage is sometimes known as asymptotic stage. Chronic stage lasts around a decade for the people who are not under any antiretroviral therapy (ART). People under HIV medication may remain stable in this stage for many decades, as drug brings down viral activeness.

Among the HIV-1 infected patients, only a small proportion (5-8%) remains clinically stable for decades without even having any ART as they maintain a normal CD4+ T cells counts (≥ 500 cells/*µ*L). They are known as long-term nonprogressors (LTNP).

People without having ART eventually progresses to AIDS which is the final stage of HIV-1 infection. At this stage viral load increases, CD4+ T cell counts drop bellow 200 cells/*µ*L and thus the critically damaged immune system fails to protect the body from pathogens.

Gene co-expression network analysis is a popular technique, widely used in several articles to model dependency structure among gene expression, such as: used to investigate the changes of expression patterns across the disease progression stages [5], [6], [7], [8], [9], [10], [11], used to identify hub genes and relevant pathways [12], [13], used to distinguish cancer risk modules [14] and utilized for biomarker selection in cancer prognosis [15]. Gene co-expression analysis in different stages of HIV progression is noticed to be carried out in literatures [5], [7], [16]. However, in most of the cases the co-expression changes are examined between acute, chronic and non-progressor stages of infection by using different methodologies. It is more important to know the gradual changes of expression pattern from uninfected stage to the AIDS stage which is overlooked in most of the previous works. Discovering community structure has been extensively utilized to extract useful knowledge about the interactions and correlations within nodes inside complex networks. In gene co-expression networks, communities (also known as groups or modules) are detected based on gene co-expression relationships among the genes in such way that intra-connections among the genes within a community is high and inter-connections among the genes in distinct communities are low. Moreover, identification key immune regulatory hub genes which interacts with HIV-1 proteins and observed inside preserved and perturbed modules across the HIV-1 stages has not been carried out extensively earlier.

In this article, we have proposed a framework to analyze the preservation and perturbation characteristics of community structures in the gene co-expression networks from uninfected to acute, acute to chronic and chronic to AIDS stages of HIV–1 disease progression using different module preservation statistics. Differentially expressed genes between all the four stages are identified by using Significance Analysis of Microarray (SAM). A widely popular co-expression analysis technique, Weighted Gene Co-Expression Network Analysis (WGCNA) based framework [17] is then utilized to construct gene co-expression network separately for all different categories of samples. Co-expression modules are identified using average linkage hierarchical clustering which uses topological overlap based measure to compute distance between gene pairs [18], [19], [20]. Here, we have constructed module eigenegene network [21] among the identified modules and formed meta-modules to investigate relationship among them.

We have also utilized different community (or module) preservation statistics first introduced in [22] to identify preservation and perturbation of meta-modules between stage pairs: uninfected-acute, acute-chronic, and chronic-AIDS. We have used two composite preservation statistics: “*Z*_*summary*_” and “*MedianRank*”. The biological significance of the meta modules were also discovered through gene ontology and pathway based analysis. Moreover, to validate the preservation statistics of meta-modules we have performed a permutation test. The results confirm the preservation characteristics of identified meta-modules are statistically significant. To examine whether our method correctly predicts the preserved/non-preserved characteristics of modules, we have utilizes various classification techniques. Classification result signifies that the proposed method correctly identifies the preserved/non-preserved meta-modules from the stage specific HIV progression dataset.

Additionally, we have detected immune regulatory genes which are differentially expressed across HIV-1 stages and also interacts with HIV-1 proteins. Among them, key immune regulatory hub genes also have been identified in the associated protein-protein interaction networks (PPINs) of the differentially expressed genes (DEGs) using twelve topological and centrality analysis methods. Subsequently, we have searched for the key immune regulatory hub genes in the preserved and perturbed meta-modules across different HIV-1 stages and thus may be potential biomarkers in HIV-1 disease progression. Further biological validation of those key immune regulatory genes may reveal some important insights into the HIV-1 disease progression.

## 2 Materials and Methods

In this section, we provide the description about the dataset used in the current study along with the adopted methodology in detail. Fig. 1 shows the overall framework utilized in the article.

**Fig. 1.**
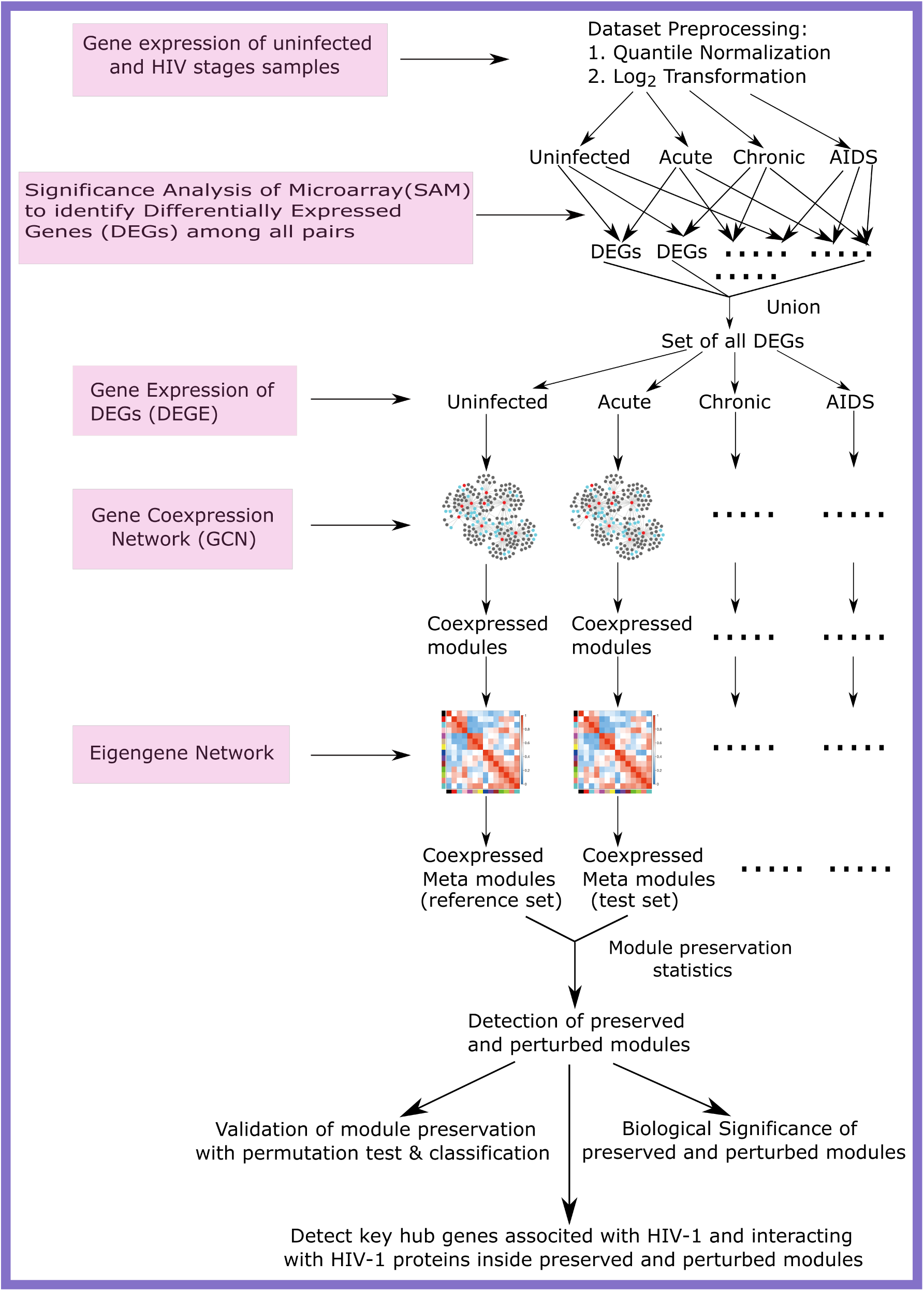
Overall framework for the present analysis.

### 2.1 Dataset Used

Qingsheng Li et al. reported stage-specific patterns of microarray gene expression during HIV-1 infection in lymphatic tissue using Affymetrix Human Genome U133 Plus 2.0 Arrays [23]. They reported a total of 52 samples measuring 54630 probe sets collected from 52 patients out of which 10 patients were unaffected, 16 in acute stage, 18 in asymptomatic (chronic) stage and 8 in AIDS stage. We have utilized this publicly available dataset comprising of four different categories of samples accessible from National Center for Biotechnology Information (NCBI) web repository to carry out our present experiment. We have used the term gene instead of probe throughout this article.

### 2.2 Dataset Preprocessing

At first, we have normalized the entire gene expression dataset using quantile normalization technique to remove different sources of systematic variations which affects the measured gene expression levels during microarray experiments. Then the normalized dataset was *log*_2_ transformed in order to easily correlate log ratio to fold change and thereby when genes are either up regulated or down regulated by factor of (say) 2, we get the same log ratio of “1” in absolute scale, “+1” for up-regulation and “–1” for down-regulation. To evaluate the differential expression of genes in order to obtain the list of up-regulated and downregulated genes across the different categories of samples, we have performed significance analysis of microarrays (SAM) [24]. SAM was carried out for each pair of categories of samples with a composite stringent filtering criterion consisting of false discovery rate (FDR) ≤ 0.025 and minimum fold change of 1.5.In SAM, a score is assigned to each gene based on its alteration in gene expression proportional to the standard deviation of gene expression across all samples of that gene. A gene is considered to be significant when its score is greater than a threshold.

Table 1 shows the statistics of differentially expressed genes (DEGs) obtained through SAM. After obtaining the DEGs for each pair of categories of samples, we have taken their union set which consists of 2829 DEGs. The expression profiles of these 2829 DEGs were used for our further analysis. From the table it can be noticed that for acute–chronic pair only 68 genes are up-regulated and 12 are down-regulated. Similarly, for chronic–AIDS, we found no up-regulated genes and only 19 down-regulated genes. The possible reason behind this statistics may be due to the fact that only a few genes play major role during the progression from acute to chronic and chronic to AIDS. Further biological validation about these genes may reveal significant findings about HIV–1 disease progression.

**TABLE 1.** Statistics of differentially expressed (DE) genes.

| Stages Compared | Upregulated | Downregulated | Total DE Genes |
| --- | --- | --- | --- |
| Uninfected – Acute | 901 | 995 | 1896 |
| Uninfected – Chronic | 782 | 776 | 1558 |
| Uninfected – AIDS | 747 | 659 | 1406 |
| Acute – Chronic | 68 | 12 | 80 |
| Acute – AIDS | 133 | 69 | 202 |
| Chronic – AIDS | 0 | 19 | 19 |
| Total (union of all DE genes) |  |  | 2829 |

Fig 2 shows the scatter plot between the average gene expression values of uninfected samples against the average gene expression values of acute samples in *log*_2_ scale. It is observed that for the acute and uninfected samples the expression values of differentially expressed up regulated genes and differentially expressed down regulated genes have the correlation values significantly greater than “1” or less than “1”, respectively. For only up regulated and down regulated genes but not differentially expressed, correlation values among the acute and uninfected samples are near about “1”.

**Fig. 2.**
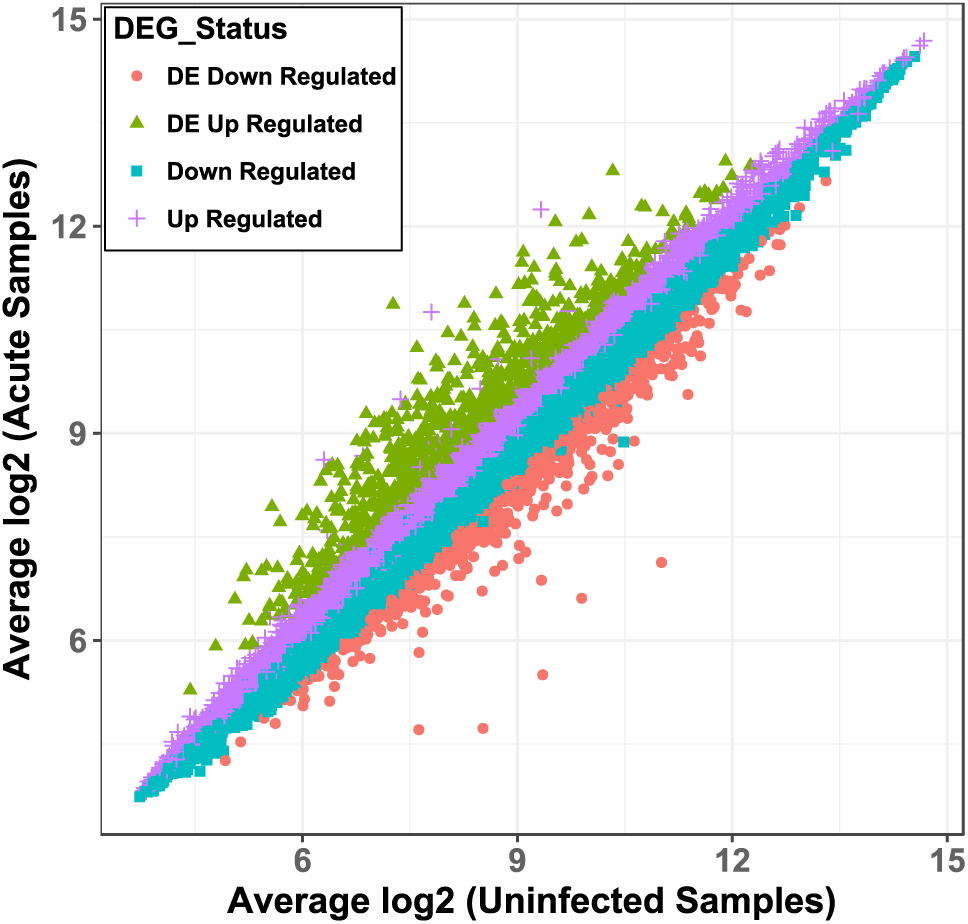
Scatter plot of average gene expression values of uninfected (control) samples vs acute samples in log_2_ scale.

### 2.3 Gene Co-expression Network Construction

We have constructed four separate gene co-expression networks (GCNs) using the normalized and *log*_2_ transformed gene expressions of the 2829 DEGs corresponding to four different categories of samples. GCNs were constructed with the help of the “weighted gene co-expression network analysis” (WGCNA) framework [17], [25]. In order to construct a GCN, at first, we have computed the pairwise gene co-expression for all DEGs using the Pearson’s correlation coefficient. Next, the adjacency matrix was constructed with the absolute values of the co-expressions. Further, the adjacencies were lifted to the soft thresholding power *λ* so that the connectivities of the resulting adjacency matrix becomes scale-free approximately (Scale-free topology criteria *R*^2^ ≥ 0.95). We have observed the *λ* values of 20, 16, 11 and 20 for control, acute, chronic and AIDS samples, respectively.

### 2.4 Identifying Co-expressed Modules

The adjacency matrix representing a GCN was converted into a topological overlap measure (TOM) based matrix in order to have a more robust similarity of the co-expression between a pair of genes considering their relative interconnectedness with every other gene within the network [18]. Then, a TOM based dissimilarity matrix was produced with which co-expression modules were detected utilizing the dynamicTreeCut algorithm that iteratively identifies stable branch size and chooses modules based on the structure of every branch (using parameters deepSplit = 2, minModulesize = 30) [26]. All the genes belonging to each of the specific modules were subsequently labeled with a unique color and genes which do not belong to any of the modules were marked with gray color.

Moreover, the expression profiles of the genes belonging to a particular module were summarized by Module Eigengene (ME) using the singular value decomposition (SVD) technique. Subsequently, we have constructed eigengene networks separately for each category of samples from which by computing the dissimilarity among the module eigengenes, meta modules were constructed utilizing the procedure mentioned in [5], [7], [21].

### 2.5 Module Preservation Statistics

In our present analysis to discover preservation and perturbation characteristics in community structure during progression of HIV–1 disease progression, we have utilized the different ‘density’, ‘connectivity’, ‘density + connectivity’ and composite module preservation statistics as stated in [22]. Module preservation measures determine the way the connectivity and the density patterns of modules in a reference network are preserved in a test network.

We have computed the different preservation statistics with 200 permutations across a reference and a test network at a time and corresponding *Z*_*statistics*_ were computed based on the results of the permutation tests [22]. Moreover, we have evaluated the value of three composite measures *Z*_*density*_, *Z*_*connectivity*_ and *Z*_*summary*_ to summarize the resultant *Z*_*statistics*_.

*Z*_*density*_ score is associated with four density measures (meanCor, meanAdj, propVarExpl, meanKME) whereas *Z*_*connectivity*_ is associated with three connectivity measures (cor.KIM, cor.KME, cor.cor). The *Z*_*summary*_ score indicates whether a module is strongly preserved (*Z*_*summary*_ ≥ 10), moderately preserved (2 ≤ *Z*_*summary*_ *<* 10), or not preserved (*Z*_*summary*_ *<* 2).

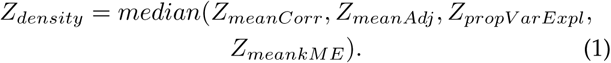

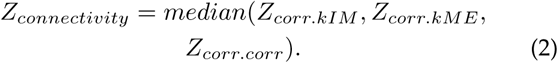

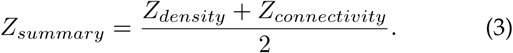

The permutation based *Z*_*statistics*_ depends on the module size. As in our case, we have modules with varying size. Thus, we have utilized another composite module preservation statistic *MedianRank* to compare relative preservation amongst modules: the lower the *MedianRank* value is, the higher is the preservation for that module. Since *MedianRank* is based on the observed preservation statistics (as compared to *Z*_*statistics*_ or other permutation test statistics), thus it is very less dependent on module size. For each density and connectivity measure, we rank the modules based on the observed score. We then define *medianRankDensity* (*mRank*_*Den*_) and *medianRankConnectivity* (*mRank*_*Con*_) analogously to *Z*_*density*_ and *Z*_*connectivity*_ from which summarized *MedianRank* was computed [6], [22].

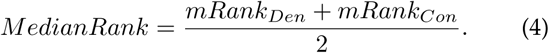

### 2.6 Identifying Key Hub Genes associated with HIV-1

When a biological system is in a state of imbalance like in disease states, the key hubs genes are typically either most up or down-regulated as they attempts to restore the system from the state of imbalance. Thus, in order to identify the key hub genes associated with HIV-1, initially, we have constituted protein-protein interactions networks (PPINs) of the strong differentially expressed genes with absolute *log*_2_ fold change greater than 2 and absolute SAM score greater than 1.5 using StringDB database. A total of 2225 genes (official gene symbols) were obtained those shown strong differential gene expression all pairs of HIV-1 stages. Thereafter, we have analyzed the obtained PPINs through twelve topological and centrality analysis methods, namely, Degree, Maximum Neighborhood Component (MNC), Density of Maximum Neighborhood Component (DMNC), Edge Percolated Component (EPC), Maximal Clique Centrality (MCC), Bottleneck, EcCentricity (ECC), Closeness, Radiality, Betweenness, Stress and Clustering Coefficient (ClCoeff) to identify the hub or essential proteins (or genes). Subsequently, we have searched for hub genes present in the preserved and non-preserved co-expression meta modules across each pair of HIV-1 stages. Further, in order to obtain hub genes across a pair of HIV-1 stages, we filtered out those genes which shown strong differential expression only within those pair of stages among union of all strong differentially expressed genes across all pairs of stages.

Moreover, we have distinguished those differentially expressed genes (or proteins) which interacts with HIV-1 proteins and also have been reported as immune regulatory genes in the Immunology Database and Analysis Portal (ImmPort), Immunome Database and Immunogenetic Related Information Source (IRIS) databases [27], [28], [29], [30], [31] in order to predict key immune regulatory genes associated with HIV-1 progression. We have also analyzed those immune regulatory genes using the above twelve topological and centrality measures which are present inside the preserved and non-preserved meta modules across the HIV-1 stages and are thus potential biomarkers in HIV-1 progression.

## 3 Results and Discussion

This section reports the results of our analysis for unveiling the intramodular and topological changes in the modular structure for pairs of different categories of samples.

### 3.1 Identification of Co-expressed Modules

We have detected co-expressed modules and subsequently meta modules within the gene co-expression networks for the control samples and each stage of HIV–1 disease using gene expression data of differentially expressed genes obtained through SAM utilizing the procedure outlined in the method section. Fig. 3(a) shows the hierarchical clustering dendrogram for gene co-expression network of the acute stage using the differentially expressed genes. We have discovered 38, 26, 17, 32 modules and 15, 15, 15, 14 meta modules for the control, acute, chronic and AIDS samples, respectively (Table 2).

**TABLE 2.** Statistics on number of modules for all samples

| Stage | Module | Meta Module | Preservation Statistics |  |  |  |
| --- | --- | --- | --- | --- | --- | --- |
|  |  |  | Compared Stage | Not Preserved | Moderately Preserved | Strongly Preserved |
| Ctrl | 38 | 15 | Acute | 5 | 9 | 1 |
|  |  |  | Chronic | 5 | 8 | 2 |
|  |  |  | AIDS | 10 | 4 | 1 |
| Acute | 26 | 15 | Chronic | 1 | 8 | 6 |
|  |  |  | AIDS | 3 | 9 | 3 |
| Chronic | 17 | 15 | AIDS | 7 | 6 | 2 |
| AIDS | 32 | 14 | Not Applicable |  |  |  |

**Fig. 3.**
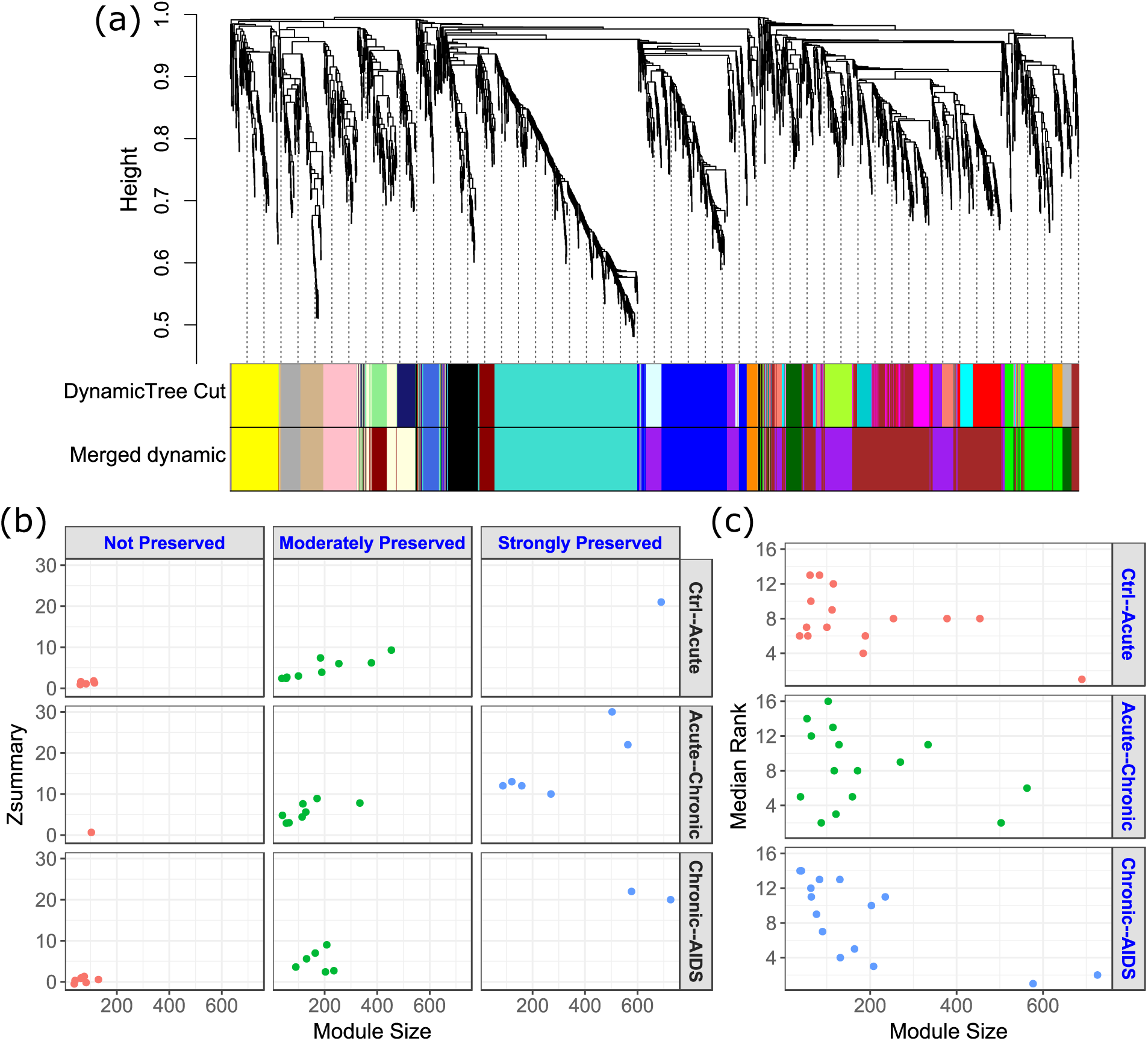
(a) figure shows the hierarchical clustering dendrogram for gene co-expression network of acute stage using the differentially expressed genes. (b) shows plots of *Z*_*summary*_ with module size of co-expression modules for control–acute, acute–chronic, chronic–AIDS stages pairs (c) figure shows scatter plots of *MedianRank* vs module size of co-expressed modules for the control– acute, acute–chronic, chronic–AIDS stages pairs.

### 3.2 Preserved Modules in Pair of Stages

In our present analysis, we have computed the module preservation statistics across the meta module sets of the different categories of samples. In particular, we have assumed co-expression meta modules of one category of samples as reference dataset and the co-expression meta modules of another category of samples as test dataset. This procedure has been performed for three pairs: uninfected (reference set) with acute (test set), acute (reference set) with chronic (test set) and chronic (reference set) with AIDS (test set) to characterize the changes in community structure in the gene co-expression network during progression of the HIV– 1 disease.

We have utilized the principle in [26]. The value of *Z*_*summary*_ above 10 denotes strongly preserved module, less than 2 represents non-preserved module while the score between 2 and 10 denotes moderately preserved module. We have sketched *Z*_*summary*_ scores against the sizes of meta modules in three columns in Fig. 3(b). Column 1 expresses non-preserved meta modules, while column 2 and column 3 depict moderately preserved and strongly preserved meta modules for the pairs of samples by considering control as reference dataset in row 1, acute as reference dataset in row 2 and chronic as reference dataset in row 3.

It can be seen from Table 2 that from our analysis only 1 strongly preserved control meta module (1 out of 15 : 6.66%) whereas 9 moderately preserved meta modules and 5 non-preserved meta modules were found across the control–acute samples’ pair by taking control meta modules as reference set and acute meta modules as test set. For co-expression meta modules of acute samples, it is observed that for the pair acute–chronic, the number of strongly preserved meta modules is higher (6 out of 15: 40%) than the other pairs of samples, 8 meta modules are moderately preserved and 1 meta module is non-preserved. This may be due to the fact that expression patterns and functionalities of only a smaller number of genes are perturbed between acute and chronic stages pair. This can be further validated using the fact that only 68 genes (affymetric id) were up-regulated and only 12 genes were down-regulated between acute and chronic stages pair using SAM analysis (see Table 1). For the samples’ pair chronic–AIDS, we have obtained 2 strongly preserved meta modules (2 out of 15: 13.33%), 6 moderately preserved and 7 non-preserved meta modules.

Additionally, we have examined the module preservation statistics *MedianRank* [22] for all co-expression meta modules of control samples by taking control as reference set and acute as test set. Similarly, we have also computed the module preservation statistics of all co-expressed meta modules of acute stage by taking acute as reference set and chronic as test set and also for the chronic stage by taking chronic meta modules as reference set and AIDS meta modules as test set. Using the compound module preservation statistics *MedianRank*, low values indicate strong preservation characteristics among modules between a pair of stages.

In Fig. 3(c), we have shown a scatter plot for the *MedianRank* values of all the modules obtained from the samples pairs control-acute, acute-chronic, chronic-AIDS. First row of that plot represents the *MedianRank* values of the control meta modules by taking control as reference dataset and acute as test dataset. Second row shows the *MedianRank* scores of the acute meta modules by taking acute as reference datasets and chronic as test dataset. Similarly, third row depicts the chronic-AIDS stage pair by taking chronic as reference dataset and AIDS as test dataset. We have identified that for the control samples 73.33% (for control-acute: 11 out of 15) meta modules have *MedianRank* less than 10, although only one module namely ‘darkolivegreen’ (*MedianRank* = 1) is strongly preserved using *Z*_*summary*_. Number of meta modules having *MedianRank* less than 10 in acute stage is 9 (for acute-chronic: 9 out of 15 modules, 60%), but 6 modules namely ‘black’, ‘blue’, ‘brown’, ‘tan’, ‘turquoise’ and ‘yellow’ are strongly preserved using *Z*_*summary*_. We have also observed that for the chronic stage only 46.66% modules (for chronic-AIDS: 7 out of 15) have *MedianRank* value less than 10 which is lower than all other samples pairs, but we have seen that two modules ‘blue’ (*MedianRank* = 2) and ‘turquoise’ (*MedianRank* = 1) are strongly preserved using *Z*_*summary*_ statistics.

### 3.3 GO and Pathway Based Analysis

Tables 3, 4 show the preservation statistics along with biological significance of the non-preserved and strongly preserved meta modules, respectively for the samples pairs control–acute, acute–chronic, chronic–AIDS. We have performed gene ontology (GO) and pathway (KEGG) based analysis to discover the biological significance of the non preserved and strongly preserved meta modules. The “Database for Annotation, Visualization and Integrated Discovery (DAVID)” [32] tool has been used for performing this analysis. First column of these tables represent module name along with size of the module. Columns 2 and 3 represent the *Z*_*summary*_ score and *MedianRank* of the meta modules. Columns 4 and 5 show the gene ontology terms and associated Kyoto Encyclopedia of Genes and Genomes (KEGG) pathways for those modules, respectively.

**TABLE 3.**
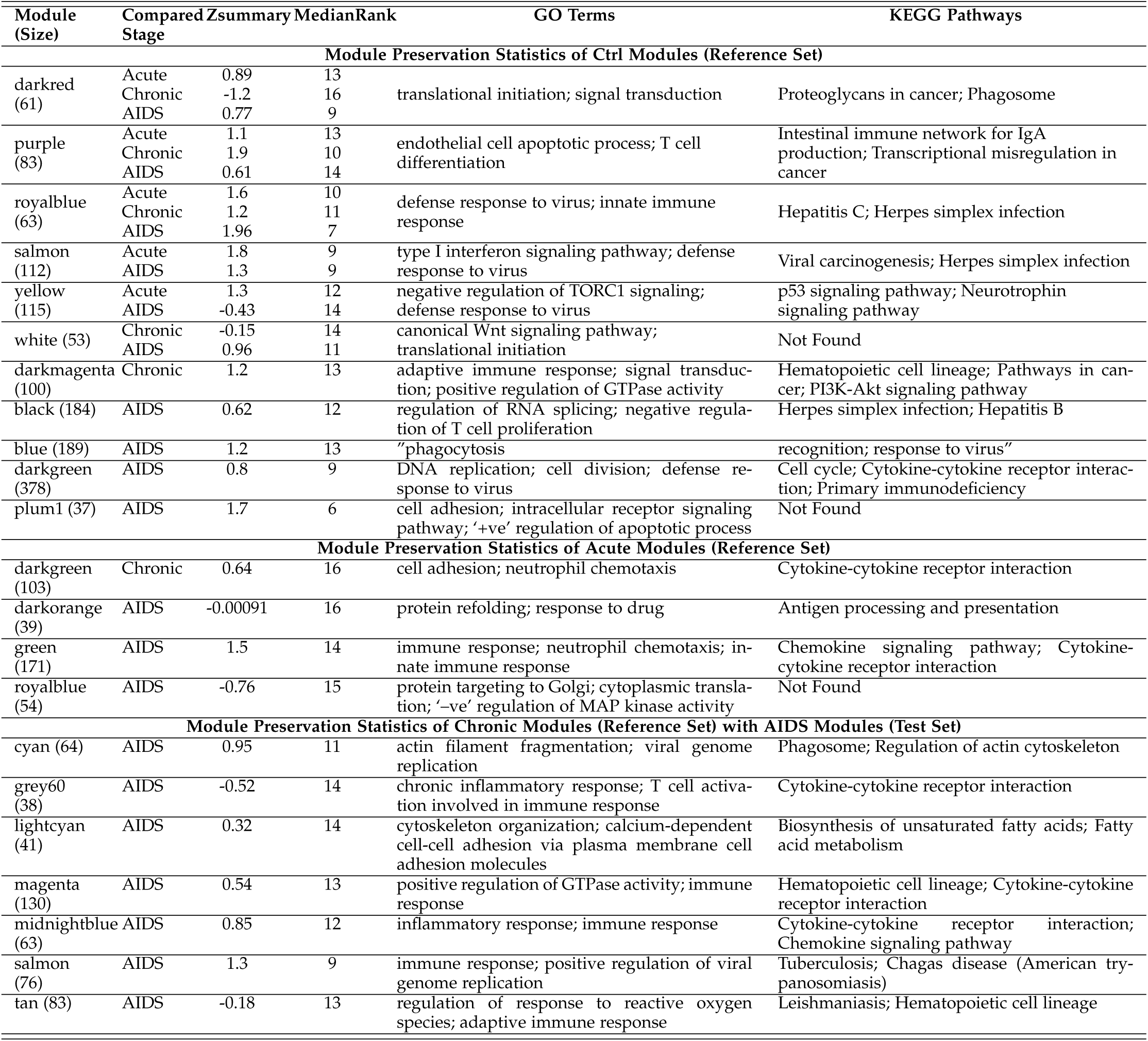
Significant gene ontology terms and pathways of non preserved modules

**TABLE 4.**
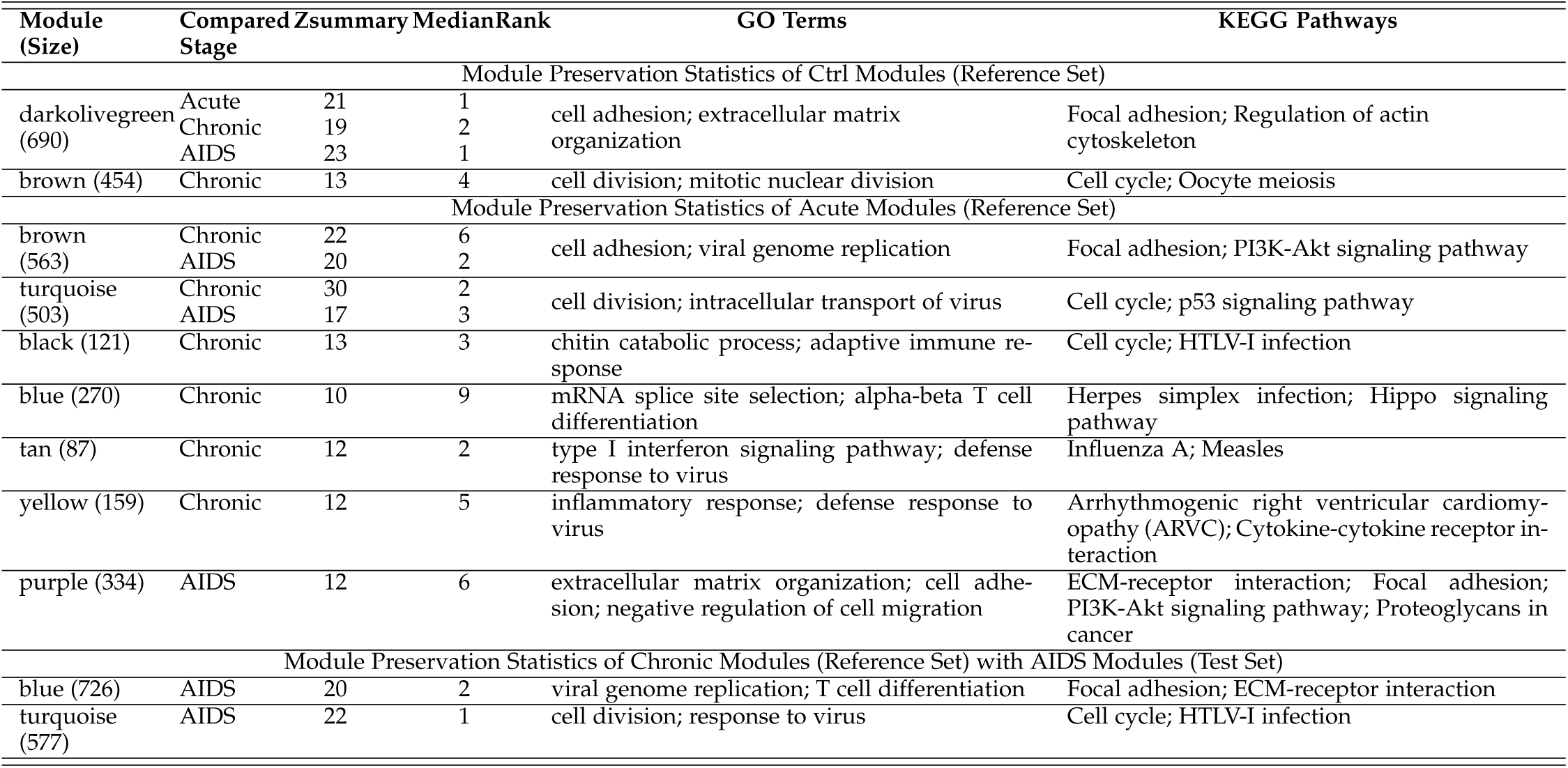
Significant gene ontology terms and pathways of strongly preserved modules

Table 3 shows the preservation statistics and biological significance of the non-preserved meta modules for the samples pairs. The ‘royalblue’ and ‘salmon’ meta modules are associated with the significant pathway as ‘herpes simplex infection’. Herpes simplex virus (HSV) is a common cause of ulcerative skin disease in both immune-compromised and immune-competent individuals. Infection can be caused by either HSV–1 or HSV–2. Epidemiological and molecular studies indicate a strong and synergistic relationship between the dual epidemics of HSV–2 and HIV–1 infection. Investigations show that HSV–2 infection enhances the risk for HIV–1 acquisition by two-three folds and HSV–2 suppression with standard prophylactic doses of HSV–2 therapy did not prevent HIV–1 acquisition [33]. The ‘darkgreen’, ‘gray60’ and ‘magenta’ meta modules are associated with the significant pathway ‘cytokine–cytokine receptor’. Cytokines are typically proteins or glycoproteins. The cytokine levels are measured in plasma in typical clinical stages. This acute seroconversion syndrome are mediated since inflammatory cytokins are characteristically exalted in plasma levels [34]. From table 3, it is quite evident that the most of the non preserved modules are enriched with some significant GO terms and significant pathways associated with the HIV–1 disease progression.

Table 4 shows the module preservation statistics along with significant gene ontology terms and pathways of strongly preserved meta modules. The ‘darkolivegreen’ meta module of control samples is associated with the significant pathway ‘regulation of actin cytoskeleton’. The cytoskeleton is made up of actin, microtubule and intermediate filament systems. As obligate intracellular pathogens, viruses depend on host cell machinery for major steps in their replication cycle and thus rely on functional interactions with cytoskeletal elements. Cytoskeleton performs an important role in the life cycle of viral pathogens whose propagation depends on mandatory intracellular steps. The actin cytoskeleton represents a significant physical barrier during virus entry associated with HIV–1 disease progression [35]. We observed ‘Influenza A’ as the pathway of ‘tan’ module of acute samples. It is known that ‘Influenza A’ virus can cause acute respiratory infection in humans and animals throughout the world. ‘Influenza virus’ infection has a great influence in HIV–positive individuals as blood cells like primary monocytes and T cells are infected by influenza virus. HIV infection is associated with poor prognosis of influenza [36]. It has been observed from the table 4 that the most of the strongly preserved modules associated with acute–chronic and chronic–AIDS are enriched with some significant GO terms and significant pathways that are strongly related to the HIV–1 disease progression. The sole strongly preserved modules associated with control samples perform basic cellular function and restrict virus replication.

### 3.4 Validation of Module Preservation Statistics using Permutation Test

To validate the preservation characteristics of obtained meta-module, we have also performed a permutation test. Corresponding to each original meta modules of control and acute samples, we have created 150 random meta modules with the same size as of original meta modules. We have also evaluated the *Z*_*summary*_ score of all the random meta modules pairwise among the control and acute samples (i.e., we have evaluated the *Z*_*summary*_ of random module 1 of control with random module 1 of acute samples, random module 2 of control with random module 2 of acute samples, and so on). Thus when considering control as reference set and acute as test set, for each original module of control samples, we have obtained 150 *Z*_*summary*_ scores applying meta module preservation procedure. Then for each original meta module of control samples, we have performed a Wilcoxon Signed Rank permutation test using 150 *Z*_*summary*_ scores of the random meta modules and *Z*_*summary*_ of the original meta module between control and acute samples. Among all the 15 original meta modules of control samples, for 12 meta modules, we obtained p-value less than 3.33*E –* 07 and for ‘plum1’ p-value is 0.0007548. For ‘orange’ and ‘white’ modules p-value are 0.2986 and 0.1365, respectively [see table 5]. This signifies that all the original modules are not random, rather they contain highly co-expressed genes and topologically significant.

**TABLE 5.**
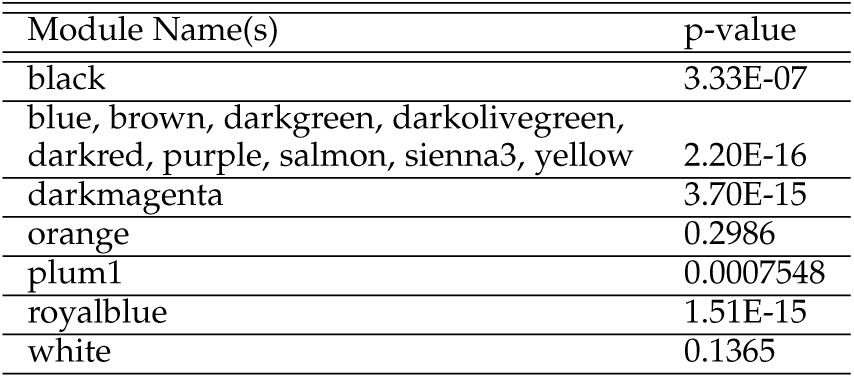
Results of permutation test for validating outcome of proposed module preservation prediction model

### 3.5 Validation of Module Preservation using Classification

To validate the efficacy of our proposed method for predicting preserved/non-preserved modules, we have performed a classification based approach. Here, we put label to the genes as ‘preserved’, ‘non-preserved’ and ‘moderately preserved’ corresponding to the modules they belong to. The expression values of each gene in four stages is treated as feature set. We have applied 7 classification techniques: k-Nearest Neighbors (kNN), Classification Tree, Stochastic Gradient Descent (SGD), Random Forest, Neural Network, Logistic Regression and AdaBoost in the dataset. We have reported 5 metrics for classification result: AUC, CA, F1, Precision and Recall in table 6. We have used 10-fold cross validation for obtaining the performance metrics. From the table, it can be noticed that most of the classification techniques provide a good classification accuracy.

**TABLE 6.**
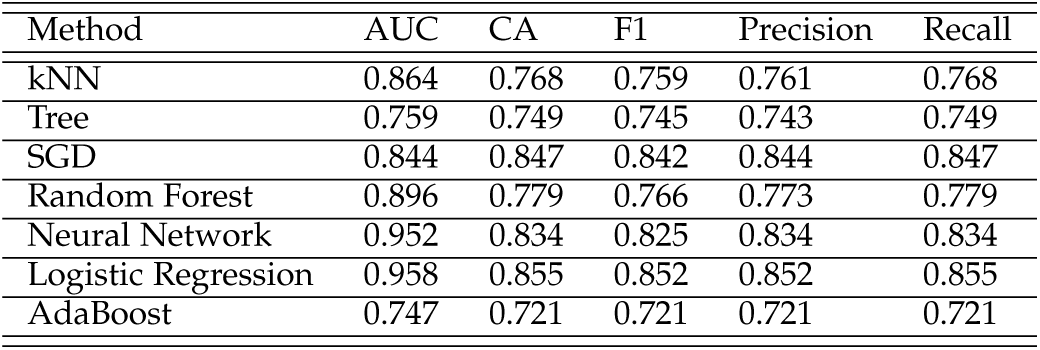
Performance of the classifiers for validating outcome of proposed module preservation prediction model

| Method | AUC | CA | F1 | Precision | Recall |
| --- | --- | --- | --- | --- | --- |
| kNN | 0.864 | 0.768 | 0.759 | 0.761 | 0.768 |
| Tree | 0.759 | 0.749 | 0.745 | 0.743 | 0.749 |
| SGD | 0.844 | 0.847 | 0.842 | 0.844 | 0.847 |
| Random Forest | 0.896 | 0.779 | 0.766 | 0.773 | 0.779 |
| Neural Network | 0.952 | 0.834 | 0.825 | 0.834 | 0.834 |
| Logistic Regression | 0.958 | 0.855 | 0.852 | 0.852 | 0.855 |
| AdaBoost | 0.747 | 0.721 | 0.721 | 0.721 | 0.721 |

### 3.6 Identifying Key Hub Genes associated with HIV-1

To identify key hub genes associated with HIV-1, we followed the procedure described in section 2.6. We obtained a PPIN of the 2225 strong differentially expressed genes (official gene symbols) that was comprises of 30283 interactions. The PPIN then was analyzed through twelve topological and centrality analysis methods to identify the hub or essential proteins (or genes) in the preserved and non-preserved co-expression meta modules across each pair of HIV-1 stages.

Subsequently, in order to obtain hub genes across a pair of HIV-1 stages, we filtered out those genes which shown strong differential expression only within those pair of stages among union of all strong differentially expressed genes across all pairs of stages. Then the key hub genes associated with HIV-1 was identified within all the hub genes by searching the genes which interacts with HIV-1 proteins [27], [28], [29], [30] and also reported as immune regulatory genes in the Immunology Database and Analysis Portal (ImmPort), Immunome Database and Immunogenetic Related Information Source (IRIS) databases [31].

There are 99 immune regulatory genes that interacts with HIV-1 proteins inside the non-preserved meta-modules in Ctrl–Acute, Ctrl-Chronic, Ctrl-AIDS stages pairs. We have also detected 45 immune regulatory genes associated with HIV-1 inside the strongly preserved modules across Ctrl– Acute, Ctrl-Chronic, Ctrl-AIDS stages pairs. Figure 4 shows the protein-protein interaction network of immune regulatory genes which interacts with HIV-1 proteins, significantly differentially expressed in HIV-1 and present inside the non-preserved and strongly preserved meta-modules in Ctrl samples across Ctrl–Acute, Ctrl-Chronic, Ctrl-AIDS stages pairs. Among the 99 immune regulatory genes of the non-preserved meta modules of ctrl samples, 17 genes, viz. ‘CXCR4’, ‘CD86’, ‘ITGB1’, ‘HERC5’, ‘FCGR3A’, ‘THBS1’, ‘ERAP1’, ‘DDX60’, ‘IFIT1’, ‘TXNIP’, ‘HSPA1A’, ‘TRAF5’, ‘FGF7’, ‘IFIT3’, ‘OAS3’, ‘C1QB’, ‘CD5L’ are present inside the ‘darkred’, ‘purple’, ‘royalblue’ meta modules which are non-preserved across all the other stages (i.e., acute, chronic, AIDS) and are thus likely to play critical roles in HIV-1 disease progression. Now on-wards, we will refer to these 17 genes as ‘Key Immune Gene Set 1’.

**Fig. 4.**
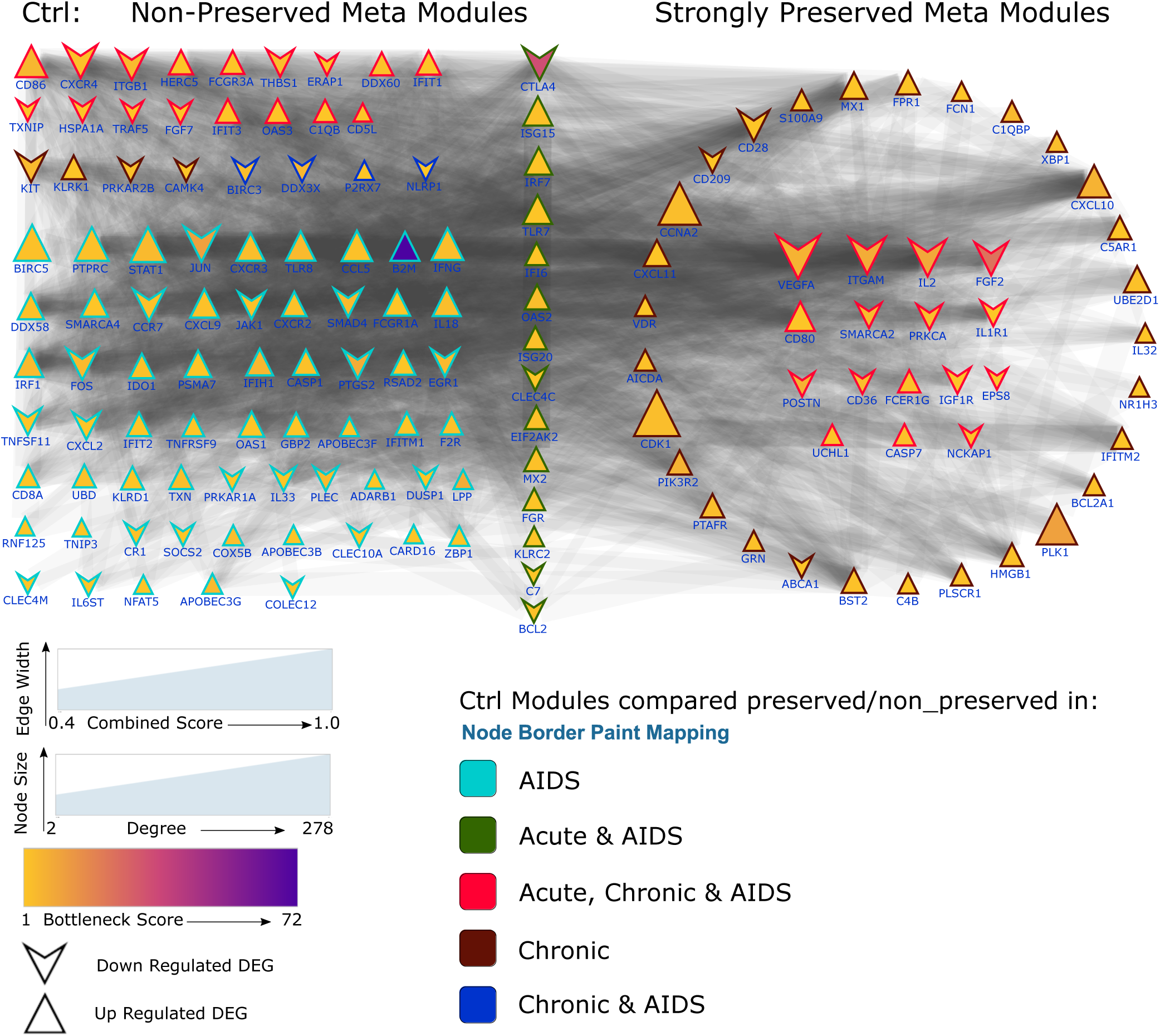
Figure shows the protein-protein interaction network of immune regulatory genes which interacts with HIV-1 proteins, significantly differentially expressed in HIV-1 and present inside the preserved and non-preserved meta-modules in ctrl samples across Ctrl–Acute, Ctrl-Chronic, Ctrl-AIDS stages pairs.

It has been also discovered that among those 99 immune regulatory genes within the non-preserved meta-modules, the genes ‘BIRC5’, ‘STAT1’, ‘PTPRC’, ‘JUN’, ‘IFNG’, ‘CD86’, ‘CTLA4’, ‘CXCR4’, ‘CCL5’, ‘TLR8’, ‘CCR7’, ‘FOS’, ‘ISG15’, ‘TLR7’, ‘B2M’, ‘CXCL9’, ‘CXCR3’, ‘IRF1’, ‘IL18’, ‘ITGB1’, ‘KIT’ have extremely high connection (degree ≥ 100) within the whole PPIN with all the 2225 strong differentially expressed genes. We will refer to these 21 genes as ‘Key Immune Hub Gene Set 2’, now onwards. These immune regulatory genes have also extremely high scores for all the 12 centrality measures viz. Degree, Maximum Neighborhood Component (MNC), Density of Maximum Neighborhood Component (DMNC), Edge Percolated Component (EPC), Maximal Clique Centrality (MCC), Bottleneck, EcCentricity (ECC), Closeness, Radiality, Betweenness, Stress and Clustering Coefficient (ClCoeff) and thus key hub or essential proteins (or genes) and potential biomarkers associated with HIV-1. Among these ‘Key Immune Hub Gene Set 2’, ‘B2M’, ‘CTLA4’, ‘JUN’ are the top 3 genes according to bottleneck centrality measures. Genes with high bottleneck centrality scores are key connector proteins with incredible functional and dynamic properties and are more likely to be essential proteins [37]. Thus all the immune regulatory genes inside the ‘Key Immune Gene Set 1’, ‘Key Immune Hub Gene Set 2’ including ‘B2M’, ‘CTLA4’, ‘JUN’ which also interacts HIV-1 proteins are likely to be essential bio-markers for HIV-1 disease progression.

Figure 5 shows the interactions among the HIV-1 proteins and the top 50 immune regulatory genes based on degree centrality measure. Among the the genes inside the ‘Key Immune Hub Gene Set 2’, ‘IFNG’, ‘CXCR4’, ‘JUN’, ‘CCL5’, ‘STAT1’, ‘FOS’, ‘CD86’, ‘CCR7’, ‘ISG15’, ‘ITGB1’ are the top 10 genes having high interactions with lots of HIV-1 proteins. Thus, these 10 immune regulatory genes have highly probable potential biomarkers in HIV-1 progression. We have also observed that the HIV-1 proteins ‘Envelope surface glycoprotein gp120’, ‘Tat’, ‘Nef’, ‘Vpr’, ‘Pr55(Gag)’ have high interactions with several immune regulatory genes which are significantly differentially expressed in HIV-1 disease progression.

**Fig. 5.**
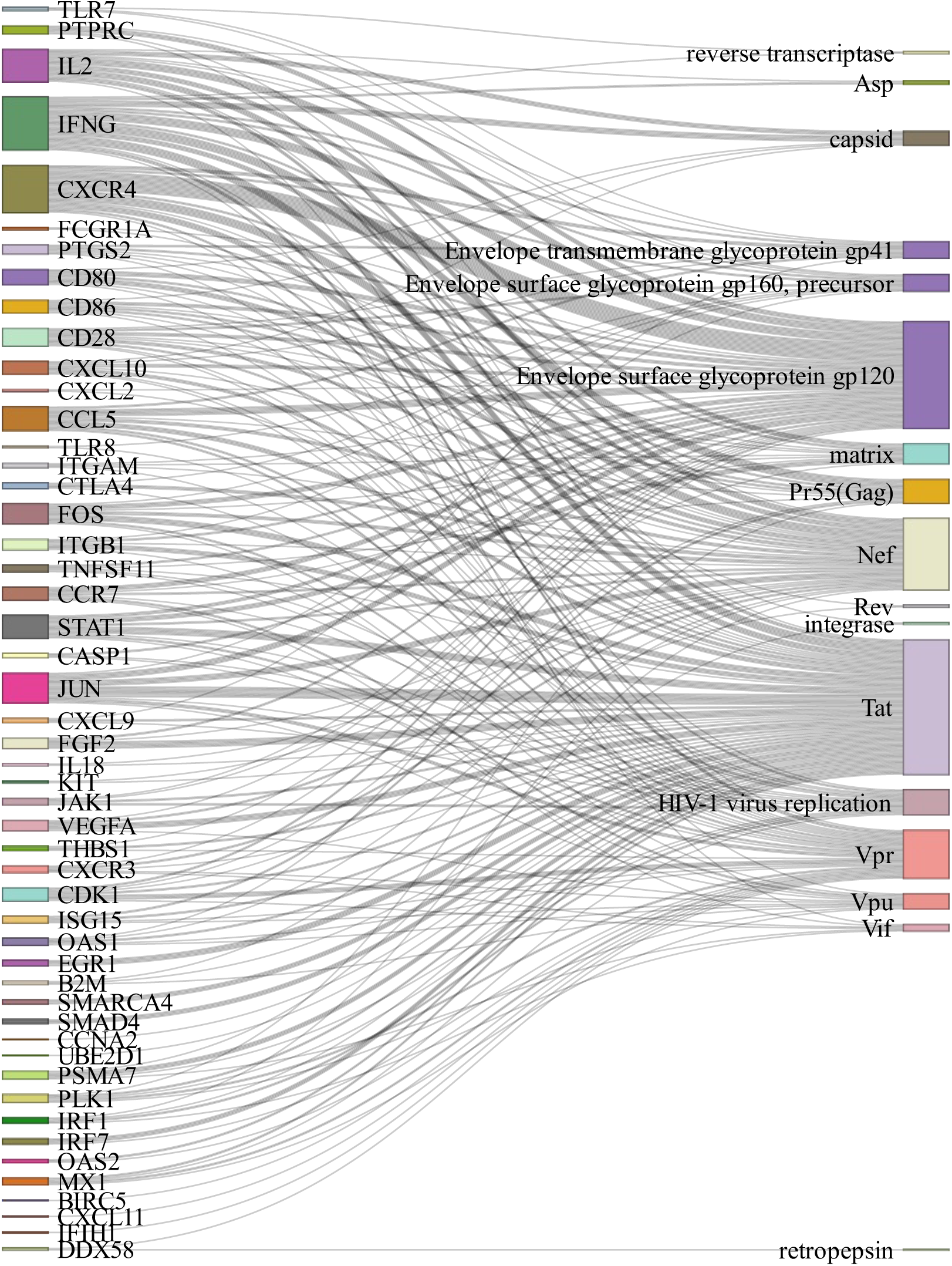
Figure shows the interactions among the HIV-1 proteins with the top 50 immune regulatory genes based on degree centrality measure.

**Fig. 6.**
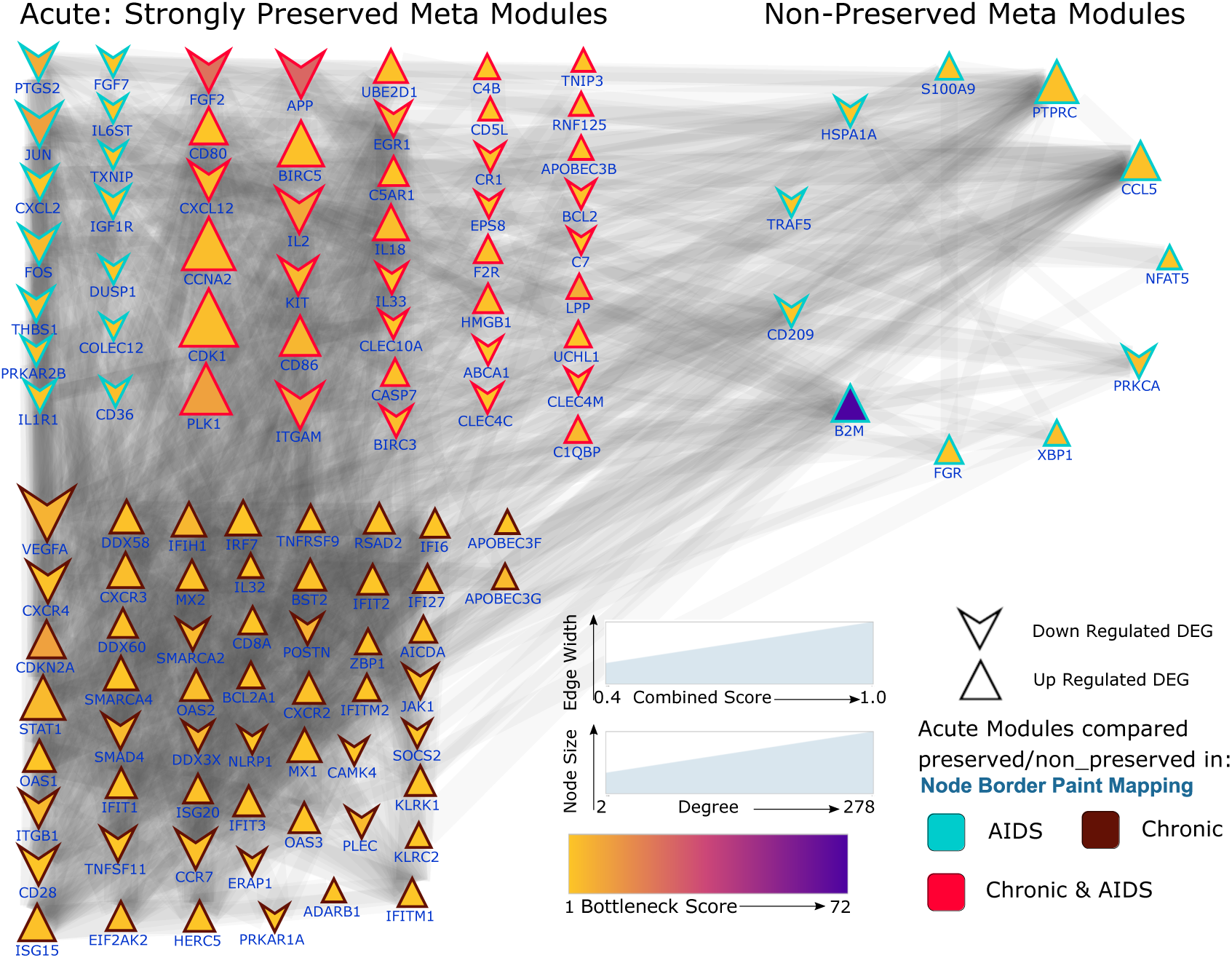
Figure shows the protein-protein interaction network of immune regulatory genes which interacts with HIV-1 proteins, significantly differentially expressed in HIV-1 and present inside the preserved and non-preserved meta-modules in acute samples across Acute-Chronic, Acute-AIDS stages pairs.

**Fig. 7.**
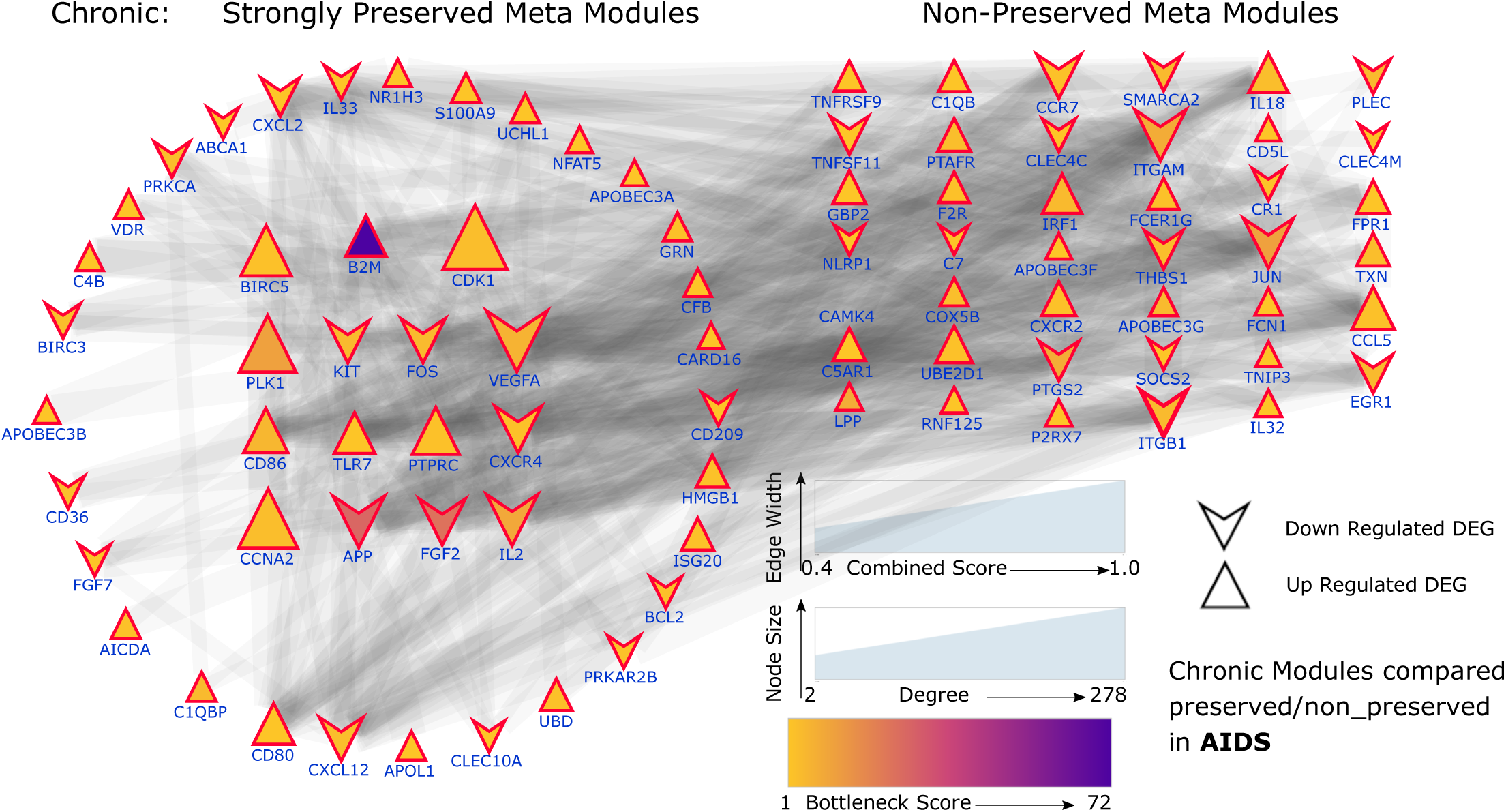
Figure shows the protein-protein interaction network of immune regulatory genes which interacts with HIV-1 proteins, significantly differentially expressed in HIV-1 and present inside the preserved and non-preserved meta-modules in chronic samples across Chronic-AIDS stages pair.

Among the 45 immune regulatory genes which are significantly differentially expressed, interacts with HIV-1 proteins and found inside the strongly preserved meta-modules across Ctrl–Acute, Ctrl-Chronic, Ctrl-AIDS stages pairs, ‘CDK1’, ‘CCNA2’, ‘VEGFA’, ‘PLK1’, ‘IL2’, ‘ITGAM’, ‘CXCL10’, ‘FGF2’, ‘CD80’, ‘CD28’ are the top 10 immune regulatory genes according to degree centrality measures (see Figure 4) and also have extremely high scores associated with all the 12 centrality measures. We will refer to these 10 genes as ‘Key Hub Gene Set 3’, now onwards. Among them ‘IL2’, ‘CDK1’, ‘VEGFA’, ‘PLK1’, ‘CXCL10’, ‘CD80’, ‘CD28’, ‘FGF2’ interacts with several HIV-1 proteins (see Figure 5).

The protein-protein interaction network of immune regulatory genes which interacts with HIV-1 proteins, significantly differentially expressed in HIV-1 and present inside the preserved and non-preserved meta-modules in acute samples across Acute-Chronic, Acute-AIDS stages pairs are shown in 6. It has been interesting fact to observe that most of the immune regulatory genes which are present inside the non-preserved meta-modules of Ctrl-Acute stage pair but not present inside the strongly preserved meta-modules across Acute-AIDS. The names of those genes are ‘B2M’, ‘PTPRC’, ‘CCL5’, ‘NFAT5’, ‘FGR’, ‘TRAF5’ and ‘HSPA1A’ and further investigation using *in-vitro* experiments may reveal some important insights about their roles in HIV-1 disease progression.

The protein-protein interaction network of immune regulatory genes which interacts with HIV-1 proteins, significantly differentially expressed in HIV-1 and present inside the preserved and non-preserved meta-modules in chronic samples across Chronic-AIDS stages pair are shown in 7. The immune-regulatory genes which are present inside the non-preserved meta-modules across Chronic-AIDS definitely play significant roles in AIDS and needs to be thoroughly investigated.

Further biological experiments using all these immune regulatory genes to validate our findings may reveal important knowledge about HIV-1 progression.

## 4 Conclusion

Our present analysis introduces a novel approach for studying the module preservation statistics of co-expression networks built using the microarray gene expression samples of uninfected data (or control) and three stages of HIV-1 disease progression. It comes out from our analysis of module preservation statistics (*Z*_*summary*_ score) that only one meta module is strongly preserved across the co-expression meta modules between control and acute samples pair by taking control as reference dataset and acute as test dataset, whereas most of the meta modules are either moderately preserved or non-preserved. The number of strongly preserved meta modules of acute samples when comparing the pair of categories of samples acute–chronic is more than the other pairs of categories of samples. This fact matches with lesser number of DEGs across acute–chronic stage pair (see Table 1) and with the outcomes of our *MedianRank* statistics where we observed low values of *MedianRank* for the strongly preserved meta modules of acute samples considering acute–chronic samples pair. We have also identified several key immune regulatory hub genes associated with HIV-1 which interacts with HIV-1 proteins inside the preserved and non-preserved modules.

Further biological validation on these immune regulatory genes inside the strongly preserved and non-preserved meta modules will certainly reveals some new facts about the progression characteristics of the HIV–1 disease. Additionally, one may investigate differential co-expression characteristics of genes during HIV–1 disease progression and apply different other statistical and machine learning algorithms like Hidden Markov Model (HMM), Deep Learning to predict novel HIV–1 associated genes. Apart from this, to biologically validate whether the genes within the strongly preserved meta modules are really engaged in HIV–1 disease, one may carry out experimental validation like gene knockdown, gene knockout mechanisms. A thorough analysis of the strongly preserved meta modules between a pair of categories of samples associated with HIV–1 disease may produce some unprecedented findings in the development of therapeutics for HIV–1 disease.

## Notes

### Competing Interest Statement

The authors have declared no competing interest.

## References

[1] C. D. Pilcher, J. J. Eron, S. Galvin, C. Gay, and M. S. Cohen, “Acute HIV revisited: new opportunities for treatment and prevention,” J Clin Invest, vol. 113, no. 7, pp. 937–945, Apr 2004.

[2] A. A. Okoye and L. J. Picker, “Cd4(+) t cell depletion in HIV infection: mechanisms of immunological failure,” Immunol Rev, vol. 254, no. 1, pp. 54–64, Jul 2013.

[3] J. M. Coquet, “Factoring in cd4 t cells during treatment of HIV,” Immunology And Cell Biology, vol. 95, pp. 571 EP –, May 2017.

[4] C. Chu and P. A. Selwyn, “Diagnosis and initial management of acute hiv infection.” American Family Physicians, vol. 81, no. 10, pp. 1239–1244, May 2010.

[5] S. Ray, S. M. M. Hossain, and L. Khatun, “Discovering preservation pattern from co-expression modules in progression of HIV-1 disease: An eigengene based approach,” in 2016 IEEE International Conference on Advances in Computing, Communications and Informatics, ICACCI 2016, Jaipur, India, September 21-24, 2016. USA: IEEE, 2016, pp. 814–820.

[6] S. Ray, S. M. M. Hossain, L. Khatun, and A. Mukhopadhyay, “A comprehensive analysis on preservation patterns of gene co-expression networks during Alzheimer’s disease progression,” BMC Bioinformatics, vol. 18, no. 1, p. 579, Dec 2017.

[7] S. M. M. Hossain, S. Ray, and A. Mukhopadhyay, “Preservation affinity in consensus modules among stages of HIV-1 progression,” BMC Bioinformatics, vol. 18, no. 1, p. 181, Mar 2017.

[8] S. M. M. Hossain, S. Ray, T. S. Tannee, and A. Mukhopadhyay, “Analyzing prognosis characteristics of Hepatitis C using a bi-clustering based approach,” Procedia Computer Science, vol. 115, no. Supplement C, pp. 282–289, 2017.

[9] M. Ray and W. Zhang, “Analysis of alzheimer’s disease severity across brain regions by topological analysis of gene co-expression networks,” BMC Systems Biology, vol. 4, no. 1, p. 136, 2010.

[10] S. M. M. Hossain, S. Ray, and A. Mukhopadhyay, “Identification of hub genes and key modules in stomach adenocarcinoma using nsnmf-based data integration technique,” in IEEE 2019 International Conference on Information Technology (ICIT), 2019, pp. 331–336.

[11] S. M. Mosaddek Hossain, A. A. Halsana, L. Khatun, S. Ray, and Mukhopadhyay, “Discovering key transcriptomic regulators in pancreatic ductal adenocarcinoma using dirichlet process gaussian mixture model,” bioRxiv, 2020. [Online]. Available: https://www.biorxiv.org/content/early/2020/10/02/2020.10.01.322768

[12] W.-C. Chou, A.-L. Cheng, M. Brotto, and C.-Y. Chuang, “Visual gene-network analysis reveals the cancer gene co-expression in human endometrial cancer,” BMC Genomics, vol. 15, no. 1, p. 300, Apr 2014.

[13] T. Wang, X. He, X. Liu, Y. Liu, W. Zhang, Q. Huang, W. Liu, L. Xiong, R. Tan, H. Wang, and H. Zeng, “Weighted Gene Co-expression Network Analysis Identifies FKBP11 as a Key Regulator in Acute Aortic Dissection through a NF-kB Dependent Pathway,” Frontiers in Physiology, vol. 8, p. 1010, 2017.

[14] X. Jia, Z. Miao, W. Li, L. Zhang, C. Feng, Y. He, X. Bi, L. Wang, Y. Du, M. Hou, D. Hao, Y. Xiao, L. Chen, and K. Li, “Cancer-Risk Module Identification and Module-Based Disease Risk Evaluation: A Case Study on Lung Cancer,” PLOS ONE, vol. 9, no. 3, pp. 1–14, 03 2014.

[15] S. Ma, M. Shi, Y. Li, D. Yi, and B.-C. Shia, “Incorporating gene co-expression network in identification of cancer prognosis markers,” BMC Bioinformatics, vol. 11, no. 1, p. 271, May 2010.

[16] S. Ray and S. Bandyopadhyay, “Discovering condition specific topological pattern changes in coexpression network: an application to HIV-1 progression,” IEEE/ACM Transactions on Computational Biology and Bioinformatics, vol. 11, no. 4, pp. 1086–1099, Dec 2015.

[17] B. Zhang and S. Horvath, “A general framework for weighted gene co-expression network analysis.” Statistical Applications in Genetics and Molecular Biology, vol. 4, pp. 1128–1172, 2005.

[18] E. Ravasz, A. Somera, D. Mongru, Z. Oltvai, and A. Barabasi, “Hierarchical organigation of modularity in metabolic networks,” Science, vol. 297, pp. 1551–1555, 2001.

[19] A. Li and S. Horvath, “Network neighborhood analysis with the multi-node topological overlap measure.” Bioinformatics, vol. 23, no. 2, pp. 222–231, Nov 2006.

[20] A. M. Yip and S. Horvath, “Gene network interconnectedness and the generalized topological overlap measure.” BMC Bioinformatics, vol. 8, no. 22, Jan 2007.

[21] P. Langfelder and S. Horvath, “Eigengene networks for studying the relationships between co-expression modules,” BMC Systems Biology, vol. 1, no. 54, Nov 2007.

[22] P. Langfelder, R. Luo, M. Oldham, and S. Horvath, “Is my network module preserved and reproducible?” PLoS Comput Biology, vol. 7, no. 1, p. e1001057, Jan 2011.

[23] Q. Li, A. J. Smith, T. W. Schacker, J. V. Carlis, L. Duan, C. S. Reilly, and A. T. Haase, “Microarray analysis of lymphatic tissue reveals stage-specific, gene expression signatures in hiv-1 infection,” J Immunol, vol. 183, no. 3, pp. 1975–1982, Aug 2009.

[24] V. G. Tusher, R. Tibshirani, and G. Chu, “Significance analysis of microarrays applied to the ionizing radiation response,” Proceedings of the National Academy of Sciences, vol. 98, no. 9, pp. 5116–5121, 2001.

[25] P. Langfelder and S. Horvath, “WGCNA: an R package for weighted correlation network analysis,” BMC Bioinformatics, vol. 9, no. 559, Dec 2008.

[26] P. Langfelder, B. Zhang, and S. Horvath, “Defining clusters from a hierarchical cluster tree: the dynamic tree cut package for R,” Bioinformatics, vol. 24, pp. 719–720, 2008.

[27] D. Ako-Adjei, W. Fu, C. Wallin, K. S. Katz, G. Song, D. Darji, J. R. Brister, R. G. Ptak, and K. D. Pruitt, “HIV-1, human interaction database: current status and new features,” Nucleic Acids Research, vol. 43, no. D1, pp. D566–D570, 11 2014.

[28] J. W. Pinney, J. E. Dickerson, W. Fu, B. E. Sanders-Beer, R. G. Ptak, and D. L. Robertson, “Hivâeur”host interactions: a map of viral perturbation of the host system,” AIDS, vol. 23, no. 5, 2009.

[29] W. Fu, B. E. Sanders-Beer, K. S. Katz, D. R. Maglott, K. D. Pruitt, and R. G. Ptak, “Human immunodeficiency virus type 1, human protein interaction database at ncbi,” Nucleic Acids Research, vol. 37, pp. D417–D422, 10 2008.

[30] R. G. Ptak, W. Fu, B. E. Sanders-Beer, J. E. Dickerson, J. W. Pinney, D. L. Robertson, M. N. Rozanov, K. S. Katz, D. R. Maglott, K. D. Pruitt, and C. W. Dieffenbach, “Short communication: Cataloguing the hiv type 1 human protein interaction network,” AIDS Research and Human Retroviruses, vol. 24, no. 12, pp. 1497–1502, 2008, pMID: 19025396. [Online]. Available: https://doi.org/10.1089/aid.2008.0113

[31] K. Breuer, A. Foroushani, M. Laird, C. Chen, A. Sribnaia, R. Lo, G. Winsor, R. Hancock, F. Brinkman, and D. Lynn2, “InnateDB: systems biology of innate immunity and beyond–recent updates and continuing curation.” Antiviral Res, vol. 41, pp. D1228–D1233, 2013.

[32] D. Huang, B. Sherman, Q. Tan, J. Collins, W. Alvord, J. Roayaei, R. Stephens, M. Baseler, H. Lane, and R. Lempicki, “The david gene functional classification tool: a novel biological module-centric algorithm to functionally analyze large gene lists,” Genome Biology, vol. 8, no. 9, p. R183, 2007.

[33] R. V. Barnabas and C. Celum, “Infectious co-factors in HIV-1 transmission herpes simplex virus type-2 and HIV-1: New insights and interventions,” Curr HIV Res, vol. 10, no. 3, pp. 228–237, Apr 2012. [Online]. Available: http://www.ncbi.nlm.nih.gov/pmc/articles/PMC3563330/

[34] M. L. Freeman, C. L. Shive, T. P. Nguyen, S.-A. Younes, S. Panigrahi, and M. M. Lederman, “Cytokines and t-cell homeostasis in HIV infection,” The Journal of Infectious Diseases, vol. 214, no. suppl 2, pp. S51–S57, Oct 2016. [Online]. Available: http://dx.doi.org/10.1093/infdis/jiw287

[35] B. Stolp and O. T. Fackler, “How HIV takes advantage of the cytoskeleton in entry and replication,” Viruses, vol. 3, no. 4, pp. 293–311, Apr 2011. [Online]. Available: http://www.ncbi.nlm.nih.gov/pmc/articles/PMC3185699/

[36] X. Wang, J. Tan, S. Biswas, J. Zhao, K. Devadas, Z. Ye, and Hewlett, “Pandemic influenza a (h1n1) virus infection increases apoptosis and HIV-1 replication in HIV-1 infected jurkat cells,” Viruses, vol. 8, no. 2, p. 33, Feb 2016. [Online]. Available: http://www.ncbi.nlm.nih.gov/pmc/articles/PMC4776188/

[37] H. Yu, p. M. Kim, E. Sprecher, V. Trifonov, and M. Gerstein, “The importance of bottlenecks in protein networks: Correlation with gene essentiality and expression dynamics,” PLOS Computational Biology, vol. 3, no. 4, pp. 1–8, 04 2007.

